# Intracellular biotransformation and disposal mechanisms of magnetosomes in macrophages and cancer cells

**DOI:** 10.1101/2023.03.15.532722

**Authors:** L. Gandarias, A.G. Gubieda, G. Gorni, O. Mathon, L. Olivi, Ana Abad-Díaz-de-Cerio, M.L. Fdez-Gubieda, A. Muela, A. García-Prieto

**Affiliations:** Dpto. Inmunología, Microbiología y Parasitología, Universidad del País Vasco - UPV/EHU, 48940 Leioa, Spain; BL22-CLÆSS Beamline, ALBA Synchrotron, 08290 Barcelona, Spain; Institute of Optics (IO-CSIC), C/Serrano 121, 28006 Madrid, Spain; BM23 Beamline, ESRF, 38000 Granoble, France; XAFS Beamline, Elettra Sincrotrone, 34149 Trieste, Italy; Dpto. Electricidad y Electrónica, Universidad del País Vasco - UPV/EHU, 48940 Leioa, Spain; BCMaterials, Bld. Martina Casiano 3rd floor, 48940 Leioa,Spain; Dpto. Física Aplicada, Universidad del País Vasco - UPV/EHU, 48013 Bilbao, Spain

**Keywords:** magnetic nanoparticle degradation, magnetotactic bacteria, magnetosomes, biomedical applications, X-ray absorption spectroscopy

## Abstract

Magnetosomes are magnetite nanoparticles biosynthesized by magnetotactic bacteria. Given their potential clinical applications for the diagnosis and treatment of cancer, it is essential to understand what becomes of them once they are within the body. With this aim, here we have followed the intracellular long-term fate of magnetosomes in two cell types: cancer cells (A549 cell line), because they are the actual target for the therapeutic activity of the magnetosomes, and macrophages (RAW 264.7 cell line), because of their role at capturing foreign agents. We show that cells dispose of magnetosomes using three mechanisms: splitting them into daughter cells, excreting them to the surrounding environment, and degrading them yielding less or non-magnetic iron products. A deeper insight into the degradation mechanisms by means of time-resolved X-ray absorption near-edge structure (XANES) spectroscopy has allowed us to follow the intracellular biotransformation of magnetosomes by identifying and quantifying the iron species occurring during the process. In both cell types there is a first oxidation of magnetite to maghemite and then, earlier in macrophages than in cancer cells, ferrihydrite starts to appear. Given that ferrihydrite is the iron mineral phase stored in the cores of ferritin proteins, this suggests that cells use the iron released from the degradation of magnetosomes to load ferritin. Comparison of both cellular types evidences that macrophages are more efficient at disposing of magnetosomes than cancer cells, attributed to their role in degrading external debris and in iron homeostasis.

**Graphical Abstract:** 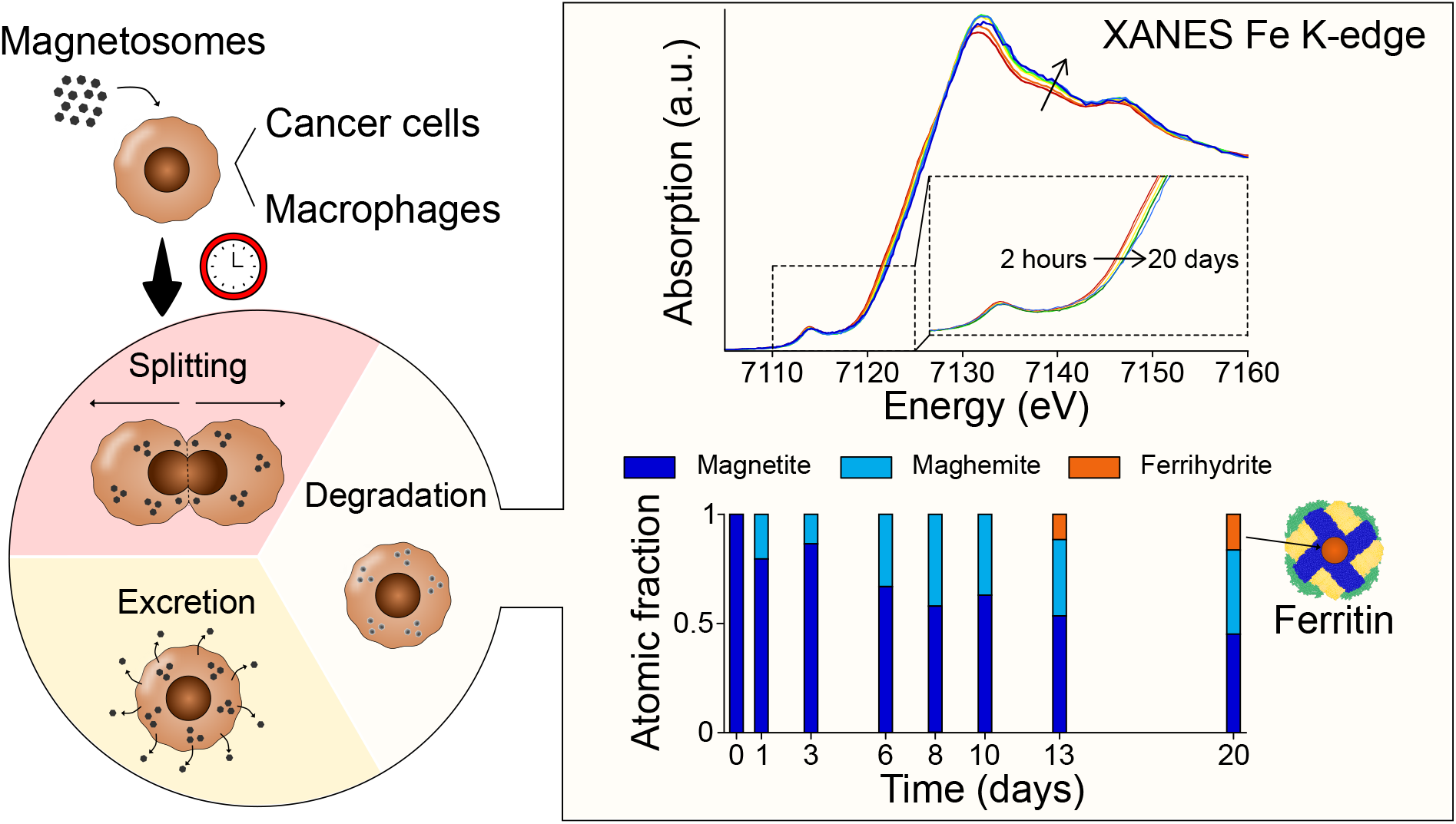

## Introduction

Magnetic nanoparticles (MNPs) have been investigated in depth over the last decades as promising biomedical agents, with the research focused on topics such as imaging, targeting, actuation, hyperthermia, and sensing, which have been extensively reviewed^1–5^. Most effort has been devoted to obtaining iron oxide MNPs due to their inherent biocompatibility and magnetic properties. Among them, magnetosomes, magnetic nanoparticles biosynthesized by magnetotactic bacteria, stand out for their remarkable properties. Magnetosomes are composed of a mineral core surrounded by a proteolipidic membrane^6–8^. The mineral core is either magnetite (Fe_3_O_4_) or greigite (Fe_3_S_4_) and presents a high chemical purity. The composition, morphology, and size of magnetosomes vary among bacterial species but are consistent within each of them, reflecting that magnetosome synthesis is under fine genetic control^6^. The size of magnetosome cores ranges from 35 to 120 nm^9^, a size window in which the particles are single magnetic domains stable at room temperature^6,8,10^. When isolated from the bacteria, the proteolipidic membrane that envelops the magnetic crystal confers the magnetosomes colloidal stability, prevents their aggregation, and facilitates their functionalization. Given these properties, the good performance of magnetosomes in biomedical applications has been proved as heating agents in magnetic hyperthermia and photothermia to treat cancer^11–13^, as drug delivery carriers^14–16^, or as contrast agents for magnetic resonance imaging (MRI)^17–20^ and for magnetic particle imaging (MPI)^21,22^. Considering the potential of magnetosomes for clinical applications, their fate after they are internalized by the body is an issue that needs to be addressed.

With this aim, here we have followed the long-term intracellular fate of magnetosomes in two cell types: cancer cells and macrophages. Cancer cells are the actual targets of the magnetosome-mediated therapies, thus, it is important to establish for how long cell internalized magnetosomes can perform their therapeutic function effectively. On the other hand, macrophages were selected because they are innate immune system cells involved in the uptake and degradation of foreign agents, such as MNPs. In fact, *in vivo* studies performed with iron oxide MNPs have evidenced that after injection, nanoparticles are predominantly accumulated in tissue-resident macrophages of liver, spleen, and kidneys^23–28^. Moreover, macrophages play a key role in iron metabolism as they are involved in iron scavenging, recycling, and storage in the body^29,30^.

The experiments were performed in A549 human lung carcinoma cells and RAW 264.7 murine macrophages, and the magnetosomes were isolated from *Magnetospirillum gryphiswaldense* MSR-1, which are composed of 40 nm sized magnetite cores. Cultures of magnetosome-loaded cells were maintained up to 20 days for A549 and up to 13 days for RAW 264.7. Using complementary techniques we followed the long term fate of magnetosomes and revealed the mechanisms that cells use to dispose of them. By means of X-ray absorption spectroscopy we were able to monitor the intracellular magnetosome degradation and to identify the iron phases occurring throughout the process.

## Materials and Methods

### Eukaryotic cell culture

A549 (DSMZ, ACC 107) and RAW 264.7 (ATCC, TIB-71) cells were cultured in RPMI-1640 medium (Sigma-Aldrich, R6504) supplemented with 2 mM L-glutamine, 10% fetal bovine serum, 100 U mL^-1^ penicillin, 100 µg mL^-1^ streptomycin, and 0.25 µg mL^-1^ amphotericin B. Cells were cultured at 37 °C in an humidified atmosphere (95 % relative humidity) and 5% CO_2_.

### Bacterial culture and magnetosome isolation

*Magnetospirillum gryphiswaldense* MSR-1 (DSM 6361) was cultured in flask standard medium (FSM) supplemented with 100 µM of Fe(III)-citrate in three-fourths 1 L bottles at 28 °C for 72 hours as described elsewere^31^. The bacteria were harvested by centrifugation (8000 g, 15 minutes, 4 °C) and the pellet was resuspended in 20 mM HEPES - 4 mM EDTA (pH 7.4) buffer. French Press (GlenMills) applying 1250 psig pressure was used for bacterial disruption. Cell lysate was sonicated to disperse cell debris from magnetosomes and these were collected and rinsed 5 times with 10 mM HEPES - 200 mM NaCl buffer (pH 7.4) and a magnetic rack. Finally, magnetosomes were collected in MilliQ water and stored at 4 °C.

### Culture of eukaryotic cells with magnetosomes

A549 and RAW 264.7 (2 *×* 10^5^ cell mL^-1^) were put in contact with 30 µg mL^-1^ magnetosomes suspended in serum free RPMI medium for 2 hours after which the culture medium was replaced for fresh medium in order to remove magnetosomes that were not attached to or inside cells. Cell cultures were maintained up to 20 days for A549 and up to 13 days for RAW 264.7. This difference in the incubation time was due to the faster growing rate of RAW 264.7 as explained in the results section. In order to maintain the whole cell population in healthy and growth promoting conditions, before confluence was achieved in the culture recipient, cells were detached and transferred to larger culture flasks with up to 500 cm^2^ surface. At certain time points (2 hours, 1, 3, 6, 8, 10, 13, and 20 days for A549 and 2 hours, 1, 3, 6, 7, 8, 9, and 13 days for RAW 264.7) cells were fixed with 2% glutaraldehyde. For each time point three replicates were assessed. The growth of cell populations and the intracellular complexity were examined using flow cytometry. To perform the rest of the measurements (i.e. ICP-AES, magnetic measurements, and XANES spectroscopy) cells were harvested by centrifugation (1200 g, 2 minutes) and the pellets were freeze dried.

### Transmission electron microscopy (TEM)

Transmission electron microscopy (TEM) images were acquired with a JEOL JEM-1400 Plus electron microscope at an accelerating voltage of 120 kV. Cells were fixed with 2% glutaraldehyde in 0.1 M Sorensen’s phosphate buffer (pH 7.4), washed several times with isoosmolar phosphate/sucrose buffer, dehydrated through an increasing ethanol concentration series, and embedded in Epon Polarbed resin in beam capsules that polymerized at 55 °C in 48 hours. A Leica UCT ultramicrotome was used to obtain ultrathin sections of 70 nm that were finally deposited onto copper grids.

### Inductively coupled plasma atomic emission spectroscopy (ICP-AES)

The concentration of iron in the cell cultures was determined by inductively coupled plasma atomic emission spectroscopy (ICP-AES) (Agilent, 5110). Freeze-dried cell samples were digested in concentrated nitric acid at 80 °C overnight.

### Magnetic measurements

Magnetic measurements of the freeze-dried cell samples were performed at room temperature on a superconducting quantum interference device magnetometer (SQUID, Quantum Design MPMS3) in DC mode.

### X-ray absorption near-edge structure (XANES)

XANES experiments were performed on the Fe K-edge (7112 eV) at the BL22 - CLÆSS beamline of the ALBA synchrotron (Spain) and at the BM23 beamline of the ESRF synchrotron (France). Samples were measured in fluorescence yield mode at low temperatures (77 K in ALBA, 10 K in ESRF) to avoid radiation damage in the cells. The monochromator used in both experiments was a double Si (111) crystal. XANES spectra were measured up to k = 9 Å ^-1^. Ref-erence samples such as horse spleen ferritin (HoSF, (Sigma-Aldrich, F7879), magnetosomes, maghemite, goethite, and others) were measured in transmission configuration.

The experimental spectra were normalized using standard procedures for background subtraction and data normalization as implemented in the free software Athena of the IFEFFIT package^32,33^. The linear combination fits to known standards were also implemented with the Athena software. The refinement was performed by minimizing the square residual in which the sum runs over the experimental points: 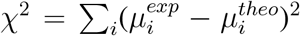 where *µ*^exp^ and *µ*^theo^ are the experimental and fitted data, respectively. The fits were considered to be reliable when they presented a *χ*^2^ *<* 0.05.

## Results and discussion

### Mechanisms of magnetosome disposal by the cells

The fate of intracellular magnetosomes was first analyzed by flow cytometry and by transmission electron microscopy (TEM). By looking into the side scattered light of cells using flow cytometry, we could follow the progression of the intracellular magnetosome content. This is because it has been proved that MNP-loaded cells have a higher value in side scattered light than unloaded cells, probably as a result of the increased intracellular complexity^34^. Figure 1a shows the progression of the side scatter in A549 and RAW 264.7. In both cases the side scatter shows a maximum after 2 hours of magnetosome uptake and decreases towards values similar to that of unloaded cells with time suggesting that cells are disposing of magnetosomes. This disposal is further confirmed by TEM in Figure 1b, where an example of A549 and RAW 264.7, 2 hours and 8 days after magnetosome uptake is displayed. It can be observed that after 2 hours magnetosomes are both attached to the cell surface and inside the cells in endosomes forming clusters, but the clusters decrease in size and number over time. Three mechanisms could be involved in magnetosome disposal: i) splitting of the magnetosome content into daughter cells due to cell division; ii) excretion of magnetosomes to the surrounding environment; iii) magnetosome degradation resulting in a reduction of their individual and cluster size.

**Figure 1:**
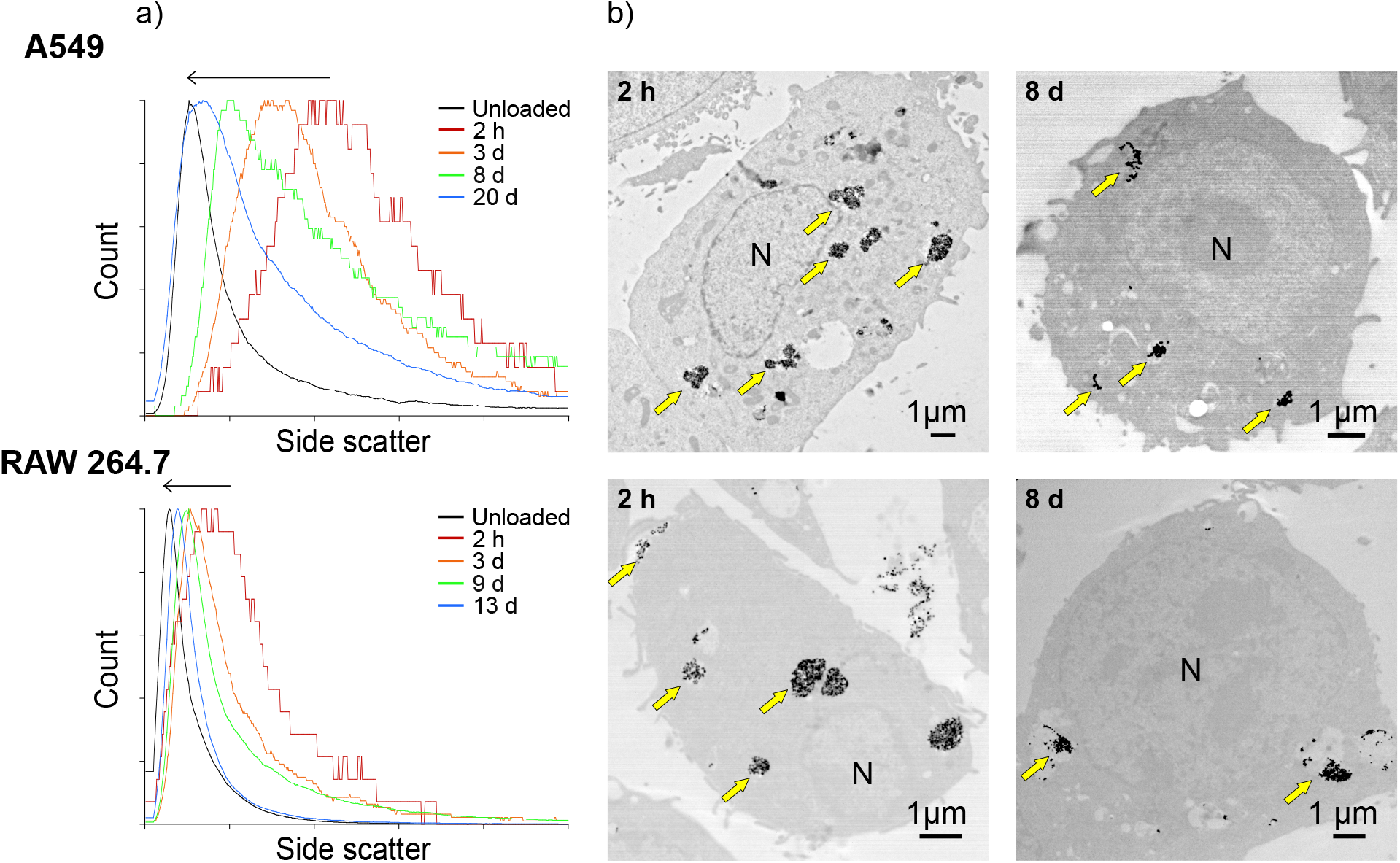
a) Histograms representing the side scattered light as a function of time in A549 and RAW 264.7. b) TEM images of A549 and RAW 264.7, 2 hours and 8 days after magnetosome uptake. Yellow arrows point at magnetosome clusters and cellular nuclei are marked N.

The first proposed mechanism (i.e. magnetosome splitting into daughter cells with cell division) is supported by the growth pattern of cell populations. As shown in Figure 2a, both A549 and RAW 264.7 populations follow an exponential growth during the experiment. The duplication time was estimated as 1.25 days for RAW 264.7 versus 3 days for A549.

**Figure 2:**
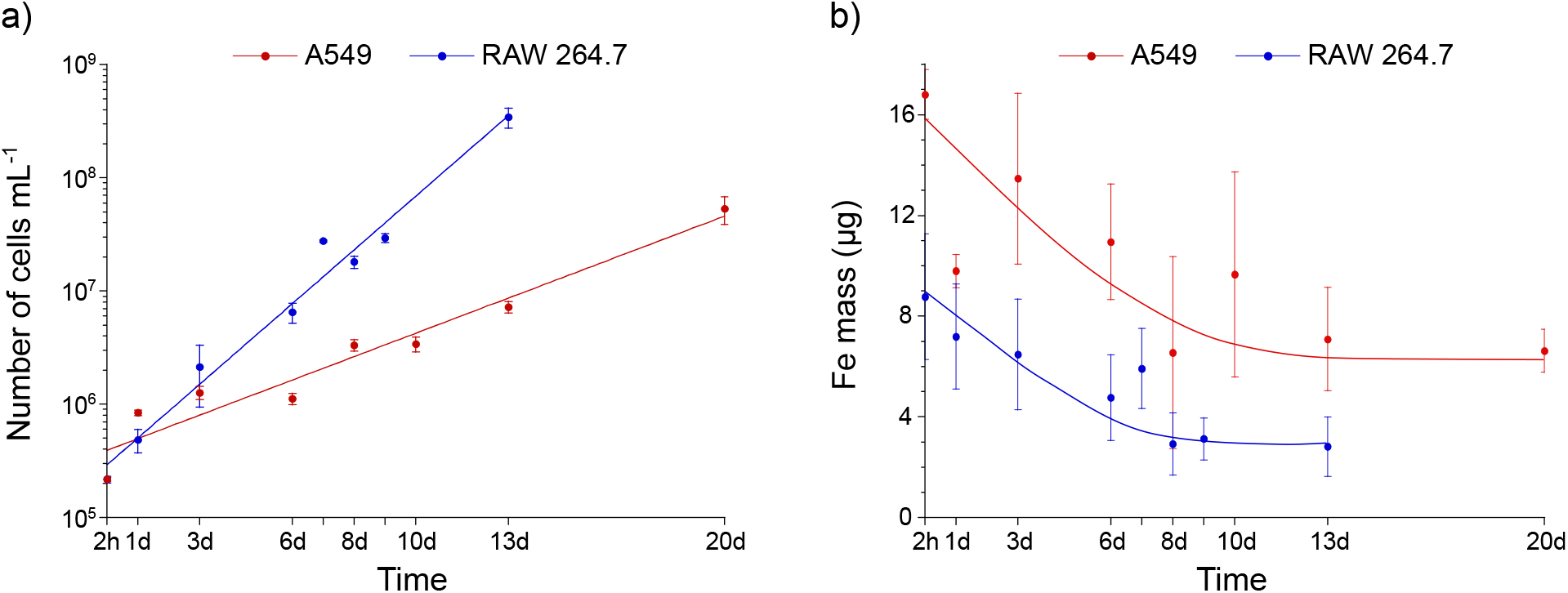
a) Progression of the number of cells as a function of time for A549 and RAW 264.7. Data represent the mean ± standard deviation, n = 3. The line is an exponential fit to the data. b) Total iron mass measured by ICP-AES of the whole cell population of A549 and RAW 264.7 along time after magnetosome uptake. Data represent the mean ± standard deviation, n = 3. The lines represent a guide for the eye.

The second proposed mechanism (i.e. excretion to the surrounding environment) is confirmed by the progression of the total iron mass of the cell populations determined by ICP-AES. As observed in Figure 2b, both cell types follow a similar trend: there is a rapid drop in intracellular iron content that stabilizes after approximately 13 days for A549 and 8 days for RAW 264.7. At the final time point measured, the decrease in the total iron mass is of ∼60% in A549 after 20 days and ∼70% in RAW 264.7 after 13 days of magnetosome uptake, indicating that macrophages are more efficient at excreting iron than cancer cells.

Further insights into the disposal of magnetosomes were obtained by following the saturation magnetic moment of cell populations using SQUID magnetometry. Figure 3a,b shows the hysteresis loops of the cell populations as a function of time. For both cell types there is a decrease of the saturation magnetic moment, *m*_*s*_, over time indicating that magnetosomes are being excreted from the cells and/or being transformed into less or non magnetic products.

**Figure 3:**
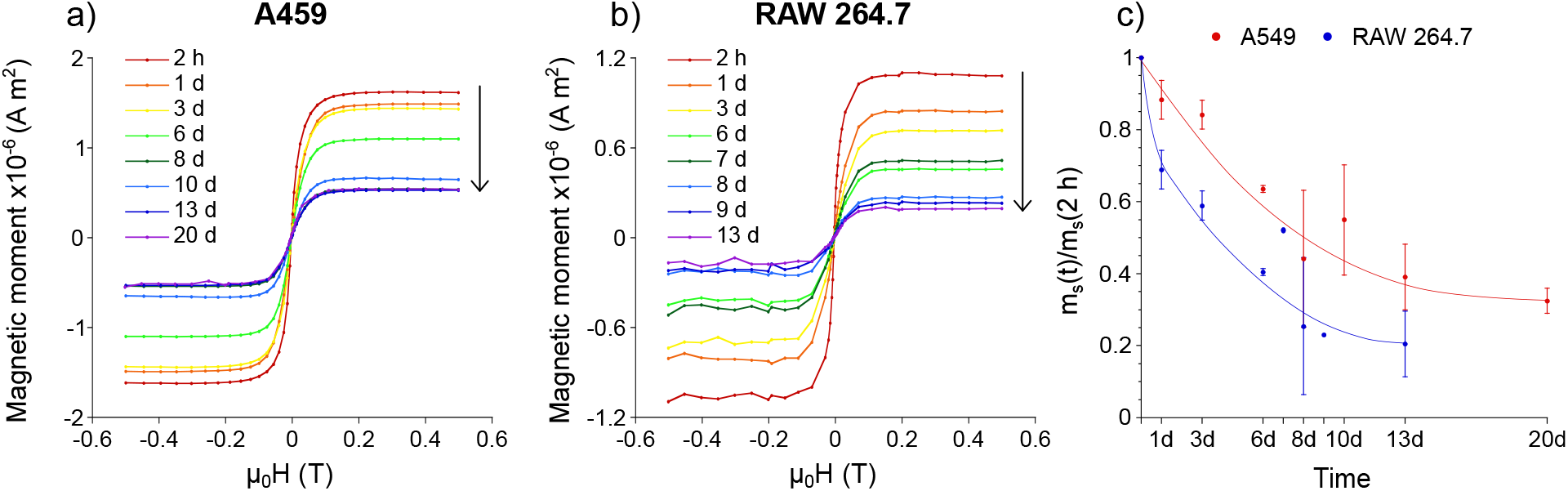
a,b) Hysteresis loops of A549 and RAW 264.7 cell populations along time. c) Ratio between the saturation magnetic moment values at a certain time point, *m*_*s*_*(t)*, and after 2 hours of magnetosome uptake, *m*_*s*_*(2 h)*. Data represent the mean ± standard deviation, n = 3. The line represents a guide for the eye.

Figure 3c shows the ratio between the saturation magnetic moment at a certain time point, *m*_*s*_(*t*), and 2 hours after magnetosome uptake by cells, *m*_*s*_(2*h*). The decrease in *m*_*s*_ was faster in RAW 264.7 as after 3 days of magnetosome uptake there was a decrease of 40% compared with the 15% in A549. Moreover, after 13 days, the *m*_*s*_ of RAW 264.7 decreased by 80% in comparison with the 70% decrease observed in A549 after 20 days. This suggests that magne-tosomes are more easily excreted and/or degraded in RAW 264.7 than in A549. Considering that magnetosome-based cancer treatments such as magnetic hyperthermia depend on the magnetic signal and often require several applications to achieve the desired effect, these results can be used as an indicator of the time progression of the treatment efficiency. Therefore, 6 days after magnetosome internalization the treatment efficiency may be reduced to ∼ 60%, which may require adjustment of the treatment parameters and/or injection of additional doses of magnetosomes. A decrease in the magnetization along time was also reported previously in human endothelial cells (HUVECs) incubated with magnetosomes^35^. However, in the same study, when working with mesenchymal stem cells, an initial decrease in *m*_*s*_ was followed by an increase to initial values in a *remagnetization* process, a phenomenon that was suggested to be stem cell specific.

### Intracellular magnetosome degradation

The third proposed mechanism of magnetosome disposal (i.e. magnetosome degradation) was followed by means of X-ray absorption near-edge structure (XANES) spectroscopy at the Fe K-edge (7112 eV). This technique allows for the specific analysis of a particular element in the sample by adjusting the energy range around its characteristic absorption edge. The XANES region provides electronic and structural information around the absorbing element, Fe in this case^36^. Using this technique we have verified intracellular magnetosome degradation and have identified the iron phases occurring during the process.

Figure 4a,d shows the Fe K-edge XANES spectra of A549 and RAW 264.7 at several time points after magnetosome uptake. Three regions of interest can be distinguished in the XANES spectra: the pre-edge, the edge, and the post-edge.

**Figure 4:**
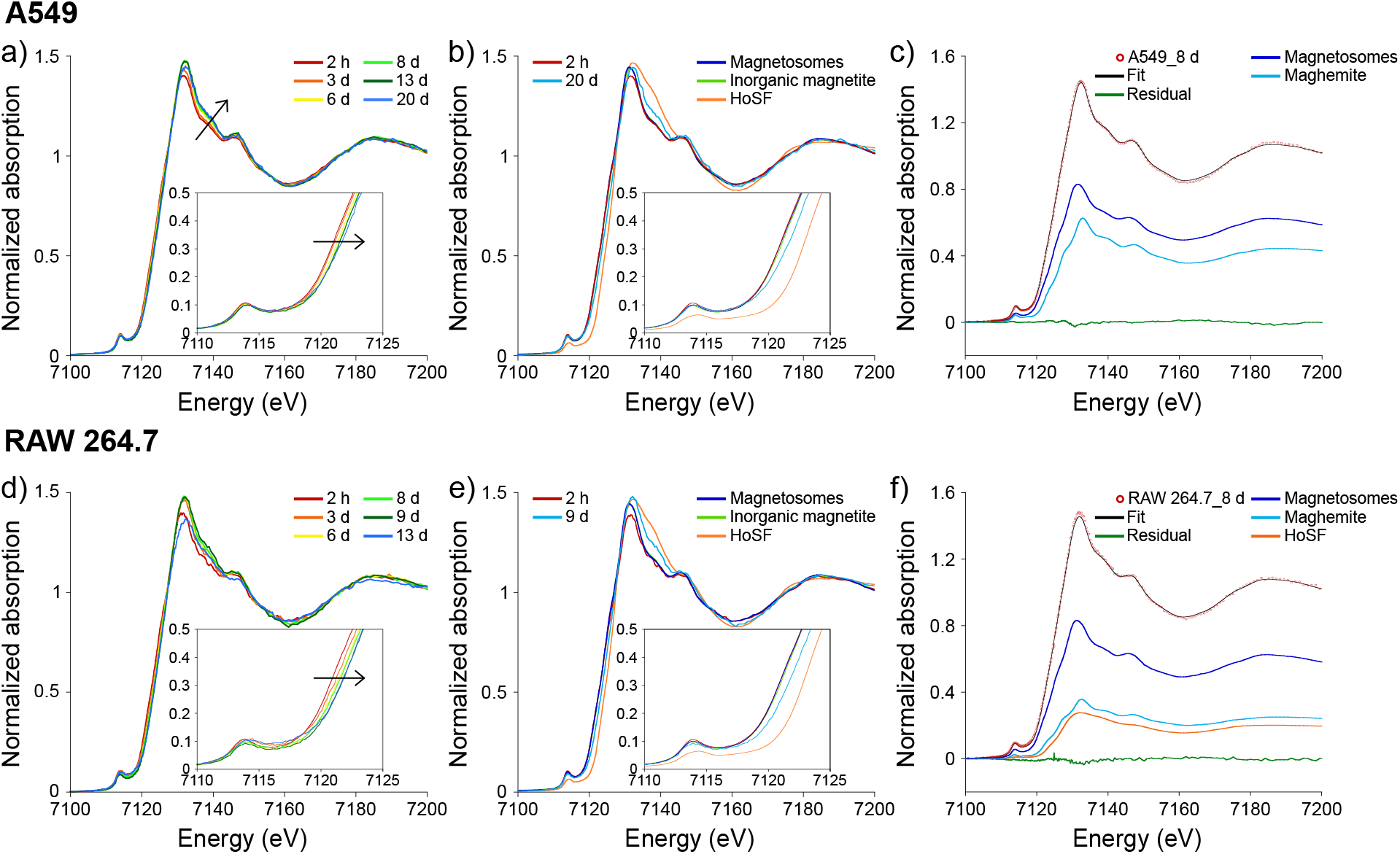
a,d) Fe K-edge XANES spectra of A549 and RAW 264.7 measured at several time points after magnetosome uptake. b,e) Fe K-edge XANES spectra of magnetosomes, inorganic magnetite (Fe_3_O_4_), and HoSF (ferrihydrite) with A549 2 hours and 20 days after magnetosome uptake and RAW 264.7 2 hours and 9 days after magnetosome uptake. c,f) Example of the linear combination fitting of A549 and RAW 264.7 XANES spectra 8 days after magnetosome uptake along with the references used for the fitting.

The pre-edge peak appears 15 - 20 eV before the main (K-edge) absorbing edge and it is usually related to the symmetry of the absorbing element. As shown in the inset of Figure 4a,d, no significant changes are observed in this region.

The edge position is a fingerprint of the oxidation state of the absorbing atom. As shown in Figure 4a,d, the edge position shifts towards higher energies over time, indicating that there is a change in the oxidation state of Fe.

The post-edge region gives information of the short range order around the absorbing atom. As observed in Figure 4a,d, there are changes at the white line (the maximum absorption just after the edge, at ∼7130 eV) and the shoulder at ∼7138 eV, indicating that the surroundings of the Fe atoms change over time.

The changes observed in the XANES spectra suggest that magnetosomes undergo intracellular degradation. As a preliminary analysis of this process, the spectra of the initial and final time points measured in this study (2 hours and 20 days for A549 and 9 days after magnetosome uptake for RAW 264.7) are compared with reference spectra in Figure 4b,e. Note that although the final time point measured for RAW 264.7 cells was after 13 days of magnetosome uptake, this spectrum deviates from the general trend and will be further explained later. As the reference spectra two iron compounds with different oxidation state are used: inorganic magnetite (Fe_3_O_4_), where iron is in a mixed oxidation state with a Fe^2+^:Fe^3+^ ratio of 1:2, and ferrihydrite, a pure Fe^3+^ compound which is the ferric oxyhydroxide mineral in the core of ferritin, a protein involved in the storage of iron in cells^37^. In this case, horse spleen ferritin (HoSF) is used as the ferrihydrite reference. The spectrum of isolated magnetosomes is also included for comparison.

The XANES spectra of A549 and RAW 264.7 2 hours after magnetosome uptake are similar to the XANES spectra of magnetosomes and inorganic magnetite, which are coincident. This confirms that magnetosomes are formed of magnetite crystals and that they remain mostly intact after 2 hours of interaction with cells. The edge positions of the final spectra, 20 days for A549 and 9 days after magnetosome uptake for RAW 264.7, are displaced to higher energies towards the Fe^3+^ edge position, but are not coincident with it. The edge position of ferrihydrite is shifted +1.58 eV with respect to magnetite. This shift is larger than the one observed in cell samples (+0.55 eV for A549 and +0.92 eV for RAW 264.7). This indicates that magnetosomes oxidize inside the cells with time but that they are not yet fully oxidized at the last time point examined. The larger shift observed for RAW 264.7 compared to A549, even in a shorter time period, could imply that they are more efficient at degrading magnetosomes.

To identify the iron phases occurring during the degradation process, a linear combination fitting of XANES spectra by using reference compounds was performed at each time point. In addition to the spectrum for magnetosomes (magnetite), which indicates the starting point, and HoSF (ferrihydrite), other references were tested: maghemite (*γ*-Fe_2_O_3_), goethite (*α*-FeO(OH)), and hematite (*α*-Fe_2_O_3_). Only three references were needed for the best fitting: magnetosomes (magnetite), maghemite, and HoSF (ferrihydrite), or goethite. An example of a linear combination fitting of A549 and RAW 264.7 8 days after magnetosome uptake is presented in Figure 4c,f. The linear combination fittings of all measured spectra along with the experimental data are represented in the Supporting Information (Figure S1).

The weight of each reference compound in the linear combination fitting represents the atomic fraction of iron in each phase (Figure 5). The degradation process follows a similar trend in both cell types. There is a first oxidation of magnetite to maghemite and then ferrihydrite starts to appear, earlier in RAW 264.7 than in A549. The appearance of ferrihydrite supports previous works that suggest that iron released from degrading magnetite nanoparticles could be locally transferred to endogenous ferritins to maintain iron homeostasis^35,38–41^. Ferritin has a protective role, as it appears to temporarily chelate labile iron that is redox-active, reducing its concentration and therefore the sensitivity of cells to oxidative stress^42,43^.

**Figure 5:**
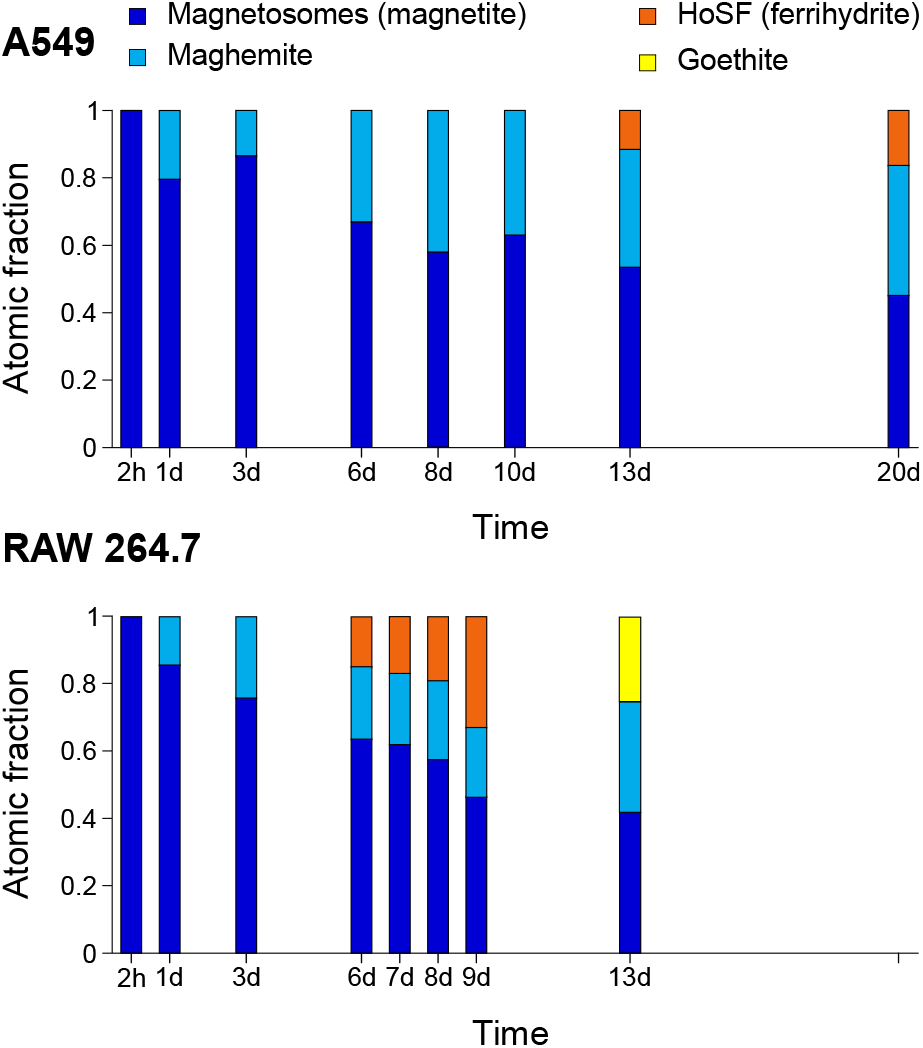
Atomic fraction of Fe in the four phases (magnetosomes (magnetite), maghemite, HoSF (ferrihydrite), and goethite) obtained from the linear combination fits of Fe K-edge XANES spectra of A549 and RAW 264.7 as a function of time after magnetosome uptake.

Despite they follow a similar trend, there is a main difference in the efficiency of both cell types in degrading magnetosomes. RAW 264.7 are more efficient than A549 as after 9 days of culture only 45% of the iron remains in the form of magnetite, compared to the 63% remaining in A549 after 10 days. Furthermore, RAW 264.7 are also more efficient in transforming the iron into ferrihydrite as after 9 days they contain 30% of the iron in this form, compared to A549 that only contain 14% of the iron in ferrihydrite after 20 days. The difference in magnetite degradation between the two cell types can be explained as RAW 264.7 are macrophages meaning that their function inside the body is to capture and degrade foreign debris whereas A549 are endothelial cells with no specific function of degradation. Moreover, macrophages play a key role in iron metabolism as they are involved in iron scavenging, recycling, and storage in the book^29,30^.

As mentioned previously, the spectrum of RAW 264.7 after 13 days of magnetosome uptake does not follow the pattern of other time points measured. For this sample the best fit was achieved with magnetosomes, maghemite, and goethite. Goethite is an iron oxyhydroxide similar to the iron mineral phase in hemosiderin^44,45^. Hemosiderin is the lysosomal degradation product of ferritin that appears under conditions of iron overload^46^. Therefore, the appearance of goethite after 13 days of magnetosome uptake could be due to the degradation of ferritin to yield hemosiderin as a consequence of an iron excess situation in the cells.

As a sign of consistency of the results, the total mass of iron obtained from ICP-AES (Figure 2b) was compared to the values estimated from the combination of the atomic fraction of iron in each phase obtained from XANES and the saturation magnetic moment, *m*_*s*_, of the hysteresis loops (Figures 3a,b and 5). This calculation is based on the fact that magnetite and maghemite are ferrimagnetic materials and contribute to *m*_*s*_, whereas ferrihydrite and goethite, being antiferromagnetic, do not contribute^47^. The details of the calculation and the results along with the total mass of iron obtained by ICP-AES are explained and represented in the Supporting Information (Figure S2). Despite some differences in the values obtained by both independent techniques, the similar overall progression of the iron content resulting from both measurements confirms the goodness of the results.

## Conclusions

In the present work we have studied the long-term fate of magnetosomes in two cell types involved in the cancer-related applications proposed for these magnetic nanoparticles: cancer cells, which are the target for the proposed treatments, and macrophages, immune cells responsible for the uptake and degradation of foreign bodies. We show that A549 human lung carcinoma cells and RAW 264.7 macrophages dispose of magnetosomes by three mechanisms: splitting into daughter cells during cell division, excretion to the surrounding environment, and intracellular degradation. The latter mechanism has been studied in depth by means of XANES spectroscopy, a technique with which we have followed the intracellular biotransformation of magnetosomes and have identified the iron phases occurring during the process. In this way, we have observed that the magnetite from magnetosomes first oxidizes to maghemite and then ferrihydrite starts to appear. The appearance of ferrihydrite suggests that cells use the magnetosome degradation products to load ferritin, a protein involved in the storage of iron in cells. We have observed that in RAW 264.7 there is a later appearance of goethite which could be related to the presence of hemosiderin, a degradation product of ferritin occurring in iron overload conditions.

The disposal mechanisms and the biotransformation process of magnetosomes follows a similar trend in both cell types analyzed. However, RAW 264.7 dispose of magnetosomes and degrade them at a faster rate than A549. For instance, when analyzing intracellular magnetosome biotransformation we have observed that ferrihydrite appears after 6 days in RAW 264.7 and after 13 days of magnetosome uptake in A549. The higher efficiency of macrophages at disposing of magnetosomes is attributed to their role in the degradation of external debris and in iron homeostasis.

This analysis represents an important step towards understanding the processing that magnetosomes undergo after being internalized by target cells, a question that needs to be addressed for the successful use of these magnetic nanoparticles as mediators in biomedical applications.

## Acknowledgements

This work was supported by the grant PID2020-115704RB-C31 funded by MCIN/AEI/10.13039 /501100011033 and by the grant IT1479-22 funded by the Basque Government. L.G. would like to thank the financial support from the grant PRE2018-083255 funded by MCIN/AEI/ 10.13039/501100011033 and by European Union NextGenerationEU/PRTR. G.G. acknowledges financial support from the grant FJC 2020-044866-I funded by MCIN/ AEI/ 10.13039/ 501100011033 and European Union NextGeneration EU/PRTR. The authors thank SGIker (UPV/EHU/ERDF, EU) for technical and human support. The authors especially thank Sergio de la Vega for his technical support, Prof. Luis Fernández Barquín and Dr. Elizabeth M. Jefremovas for the use of their SQUID magnetometer, Dr. Giuliana Aquilanti for reviewing the manuscript, and the synchrotrons ALBA (2019023274), ELETTRA (20190147), and ESRF (LS-2966) for beamtime allocation. Finally, the authors would like to extend their most sincere gratitude to Dr. Thomas Buslaps for his help and support during the experiment performed at the ESRF.

