## Supporting Information for "Intracellular biotransformation and disposal mechanisms of magnetosomes in macrophages and cancer cells"

April 21, 2023

†Current address: Bioscience and Biotechnology Institute of Aix-Marseille (BIAM), UMR7265, Aix-Marseille Université, CNRS, CEA Cadarache, 13108 Saint-Paul-lez-Durance, France.

### Linear combination fitting of the XANES spectra of cell populations

Figure S1 shows the linear combination fits of the XANES spectra of cell populations at different time points after magnetosome uptake.

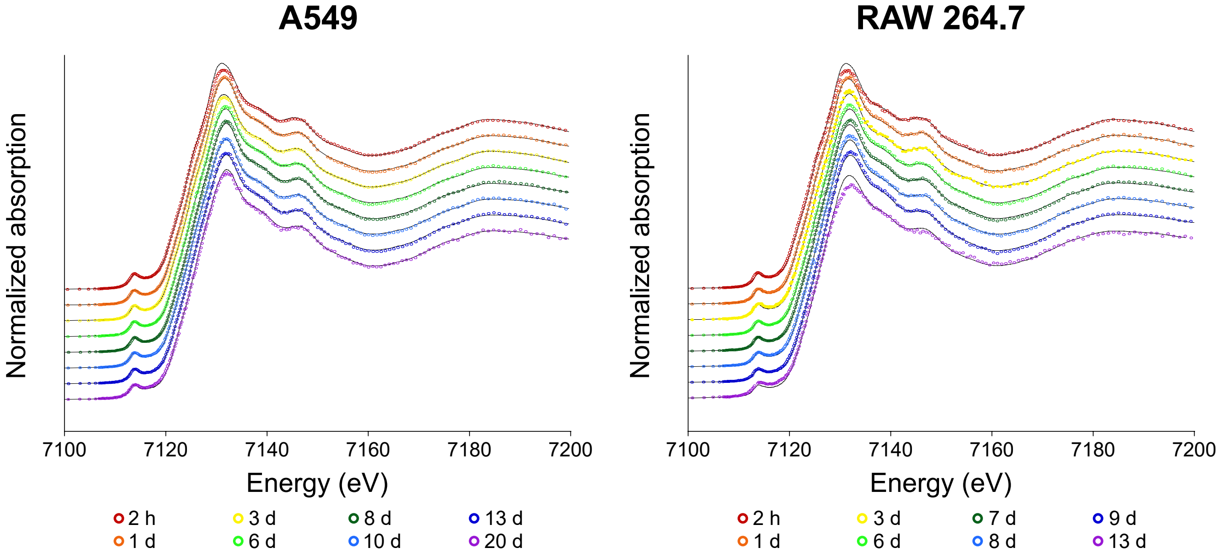


Figure S1: Fe K-edge XANES spectra of A549 and RAW 264.7 cells measured at different time points after magnetosome uptake and the corresponding linear combination fits.

### Calculation of the total mass of Fe by XANES and magnetometry

In order to verify the goodness of the results, we compared the values of the total iron mass obtained by measuring cell populations with ICP-AES and with magnetic measurements and XANES spectroscopy.

The mass of Fe in the magnetosome-loaded cells was estimated by means of the combination of the atomic fraction of Fe in each phase obtained from XANES and the saturation magnetic moment, $m_s$, of the hysteresis loops.

Considering that there are three iron phases in the samples (magnetite, maghemite, and ferrihydrite), the mass of Fe in the sample, $m_{Fe}$, is given by:

$$m_{Fe} = m_{Fe,magn} + m_{Fe,magh} + m_{Fe,fh} \tag{1}$$

From the linear combination fitting of XANES spectra, the atomic fraction of Fe in each phase is obtained: $\alpha_{magn}$, $\alpha_{magh}$, and $\alpha_{fh}$. Therefore, the Fe mass in each phase is given by:

$$\begin{aligned} m_{Fe,magn} &= \alpha_{magn} \times m_{Fe} \\ m_{Fe,magh} &= \alpha_{magh} \times m_{Fe} \\ m_{Fe,fh} &= \alpha_{fh} \times m_{Fe} \end{aligned} \tag{2}$$

$m_{Fe,magn}$ and $m_{Fe,magh}$ can be estimated from the combination of the atomic fractions obtained by XANES and the hysteresis loops because only magnetite and maghemite contribute to the saturation magnetic moment, $m_s$. Magnetite contributes with 92 A $m^2$ $kg^{-1}$ and maghemite contributes with 76 A $m^2$ $kg^{-1}$. Since the mass fraction of Fe in magnetite and maghemite is 72% and 70%, respectively, the saturation magnetic moment per mass of Fe is 127.8 A $m^2$ $kg^{-1}_{Fe}$ for magnetite and 108.6 A $m^2$ $kg^{-1}_{Fe}$ for maghemite.

From XANES, the atomic fraction of Fe in the magnetic phases (magnetite and maghemite) is given by:

$$x_{magn} = \frac{\alpha_{magn}}{\alpha_{magn} + \alpha_{magh}} \quad \text{and} \quad x_{magh} = 1 - x_{magn} = \frac{\alpha_{magh}}{\alpha_{magn} + \alpha_{magh}} \tag{3}$$

Therefore, the mass of Fe in each magnetic phase can be calculated as:

$$\begin{aligned} m_{Fe,magn} &= \frac{x_{magn}}{x_{magn} \times 127.8 + x_{magh} \times 108.6} \times m_s \\ m_{Fe,magh} &= \frac{x_{magh}}{x_{magn} \times 127.8 + x_{magh} \times 108.6} \times m_s \end{aligned} \tag{4}$$

From Equations 2 the mass of iron in ferrihydrite can be calculated as:

$$m_{Fe,fh} = \frac{\alpha_{fh}}{\alpha_{magn}} \times m_{Fe,magn} \quad \text{or} \quad m_{Fe,fh} = \frac{\alpha_{fh}}{\alpha_{magh}} \times m_{Fe,magh} \tag{5}$$

The total mass of Fe in the sample, $m_{Fe}$, can be calculated using Equation 1. The same reasoning applies when the sample contains goethite instead of ferrihydrite.

The results obtained from the calculations are observed in Figure S2.

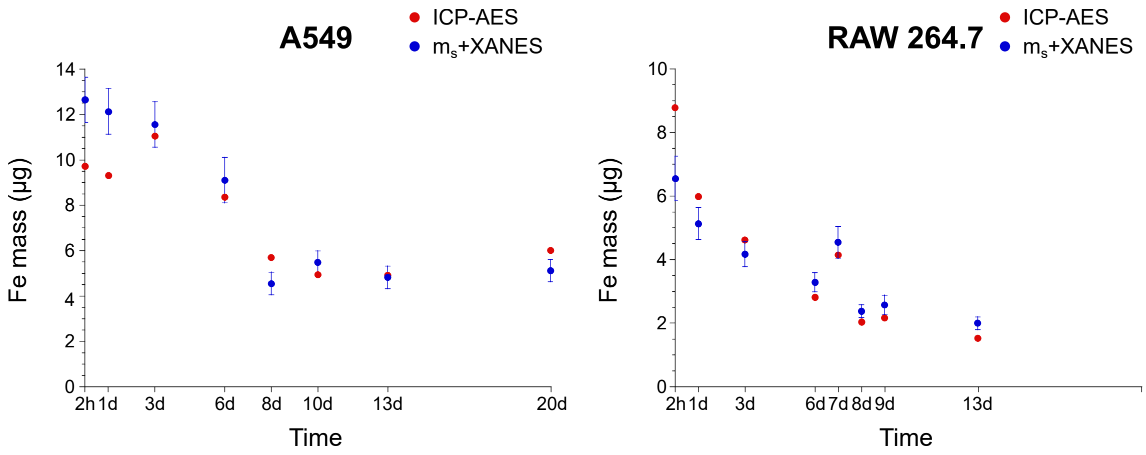


Figure S2: Mass of iron obtained by ICP-AES and estimated by the combination of the atomic fraction of Fe in each phase obtained from XANES and the saturation magnetic moment, $m_s$, of the hysteresis loops. The error of $m_{Fe}$ has been estimated by error propagation[1].
